# Nasal and gut microbiota for sows of different health status within six commercial swine farms from one swine production system

**DOI:** 10.1101/596130

**Authors:** Andréia Gonçalves Arruda, Loic Deblais, Vanessa Hale, Monique Pairis-Garcia, Vishal Srivastava, Dipak Kathayat, Anand Kumar, Gireesh Rajashekara

## Abstract

Sow culling is an essential practice in swine herds to optimize animal health and productivity; and cull sows represent a considerable proportion of the herd at any given time point. Even though recent studies have reported that the microbiome is associated with susceptibility to diseases, the microbiome in the cull sow population has not been explored. The main objective of this study was to investigate whether there were differences in abundance and diversity of microbes encountered in the gut and upper respiratory tract of sows of different health status (healthy, cull, and compromised cull sows) and different farms. Farms were visited once, 30 individual fecal and nasal swab samples were obtained per farm; and pooled across animals by health status and farm in pools of five. Genomic DNA was extracted and samples were subjected to MiSeq 16S rRNA sequencing using Illumina MiSeq. Diversity analyses were conducted using QIIME. Alpha diversity was analyzed using observed OTUs, PD Whole Tree, and Chao1; and beta diversity was assessed using weighted UniFrac. The mean number of OTUs was 3,846.97±9,078.87 and 28,747.92±14,090.50 for nasal and fecal pooled samples, respectively. Diversity of the nasal microbiota was low compared to the gut microbiota. For nasal samples, there was a difference in diversity between samples from farms 1-6 using the Chao1 metric (p = 0.0005); and weighted beta diversity values indicated clustering by health status. For fecal samples, there was no difference in diversity between compromised, cull, and healthy sows; or between samples from farms 1-6. Weighted PCoA analyses showed an influence of farm of origin on the diversity of pooled fecal samples. Finally, differences at the genus level were found in the fecal microbiota composition of sows of different health status and farm of origin; but not for nasal microbiota.

## Introduction

Culling refers to the process of removing animals from a breeding herd in order to optimize productivity and profitability, and is an essential and common practice in swine herds [1-3]. Sows, which refer to reproductively mature females, are normally culled for reasons that include but are not limited to low production efficiency, poor reproductive traits, and/ or presumed compromised immune status. Most often, the definitive underlying reason for culling is not completely elucidated. Culling rates in the U.S. average around 50% [2]. Given the logistical complications associated with pig transport, culls sows may remain on the farm for a significant period of time and represent an existing and constant subpopulation within swine herds. Even though recent work has focused on evaluating cull sows outside of the farm (i.e. auctions, sale yards etc.) [4], not much attention has been devoted to these animals as it refers to their potential role in maintenance and re-emergence of pathogens and, ultimately disease. Considering these animals are usually older, commonly immunocompromised and therefore have higher chances of previous exposure to pathogens, it has been anecdotally noted that cull sows might be a potential population from which pathogens could re-emerge within a herd.

The microbiome is defined as the microbial composition within a body site; and this large number of microbes is known to play an important role in human and animal health [5-7] and animal production [8]. The study of the swine microbiome and its impact in respiratory and systemic diseases has been an emerging study area within the past few years [9]. Previous work has explored differences between swine microbiota in farms of different health status and different countries [10]; however, a considerable challenge to date is to differentiate system or spatial effects from disease effects. Furthermore, it has been recently reported under laboratory and semi-experimental conditions that both gut and nasal microbiome are associated with susceptibility to respiratory conditions such as porcine respiratory and reproductive syndrome (PRRS) [11], porcine circovirus type 2 associated disease [11], and Glasser’s disease [10]. However, information regarding the microbiota in the cull sow population has not been explored; and to the authors’ knowledge, comparisons between the microbiota of animals from different health status that are housed within multiple commercial farms under the same management practice (e.g. natural health challenges) has not been previously reported.

The main objectives of this project were to characterize baseline gut and nasal microbiota for adult sows under commercial conditions, and to investigate whether there were differences in the abundance and diversity of microbes encountered in the gut and upper respiratory tract of sows of different health status (including healthy, cull, and compromised cull sows) and from six different farms under the same swine production system (company).

## Material and methods

### Study population and sampling protocol

This study has been approved by The Ohio State University’s Institutional Animal Care and Use Committee (Protocol #2017A00000060), as well as by the Institutional Biosafety Committee (Protocol #2017R00000041).

A cross-sectional observational study was conducted during July of 2017. The project included enrollment of adult sows distributed within six different privately owned commercial farrow-to-wean swine farms from the same production company located in the Mideast. Basic farm descriptors are shown in Table 1. One hundred and eighty animals were selected and sampled across the six farms; comprising of 60 healthy sows, 60 cull sows, and 60 compromised sows. Healthy sows were defined as adult female pigs that were part of the swine herds, either gestating, lactating, or weaned (i.e. sows that have just weaned piglets and are between weaning piglets and being bred for her next pregnancy), that had no evidence of a health condition that would compromise their permanence in the swine herd. A cull sow was defined as an adult female that was identified as a cull animal, which meant that it was to leave the farm in the next load for the culling station. For this category, a visual assessment was not enough to identify these animals as cull sows, which meant that the reasons for culling were based on decreased productive and/ or reproductive parameters. Finally, compromised cull animals were defined as sows presenting acute clinical signs of disease that could be visually assessed by the attending veterinarian. A few examples of this category included evident lameness, severe shoulder ulcers, active abortion, and poor body condition score (score 1).

**Table 1.** Farm demographics and general information

| Descriptor | Farm 1 | Farm 2 | Farm 3 | Farm 4 | Farm 5 | Farm 6 |
| --- | --- | --- | --- | --- | --- | --- |
| Date sampled | 6/26/2017 | 7/10/2017 | 7/10/2017 | 7/12/2017 | 7/12/2017 | 7/18/2017 |
| Number of sows | 1,300 | 2,700 | 1,100 | 2,400 | 5,500 | 4,000 |
| Cull sow housing type | Crates | Crates | Crates | Crates/<br>Pens | Crates/<br>Pens | Crates |
| Cull sow removal frequency | Every 2 weeks | Every 1-2 weeks | Every 2 weeks | Every 2 weeks | Weekly | Every 2 weeks |

Farms were visited once during the month of July of 2017 for collection of fecal samples and nasal swabs from all sows within a farm on the same day. A total of 30 fecal samples and 30 nasal swabs were obtained per farm (ten for each type of animal) for a total of 360 samples (180 fecal samples and 180 nasal swabs). A schematic of sampling design is presented in Fig 1. For nasal swab sampling, pigs were restrained using a snout snare, the swab was inserted into the ventral passage of the nose and left for 3 seconds and then the same procedure was conducted with the same swab for the other nasal cavity and placed in a dry tube. Following this, the fecal sample was collected by the investigators using a sterile glove using digital palpation of the rectum to remove feces which was then placed into sterile bags.

**Fig 1.**
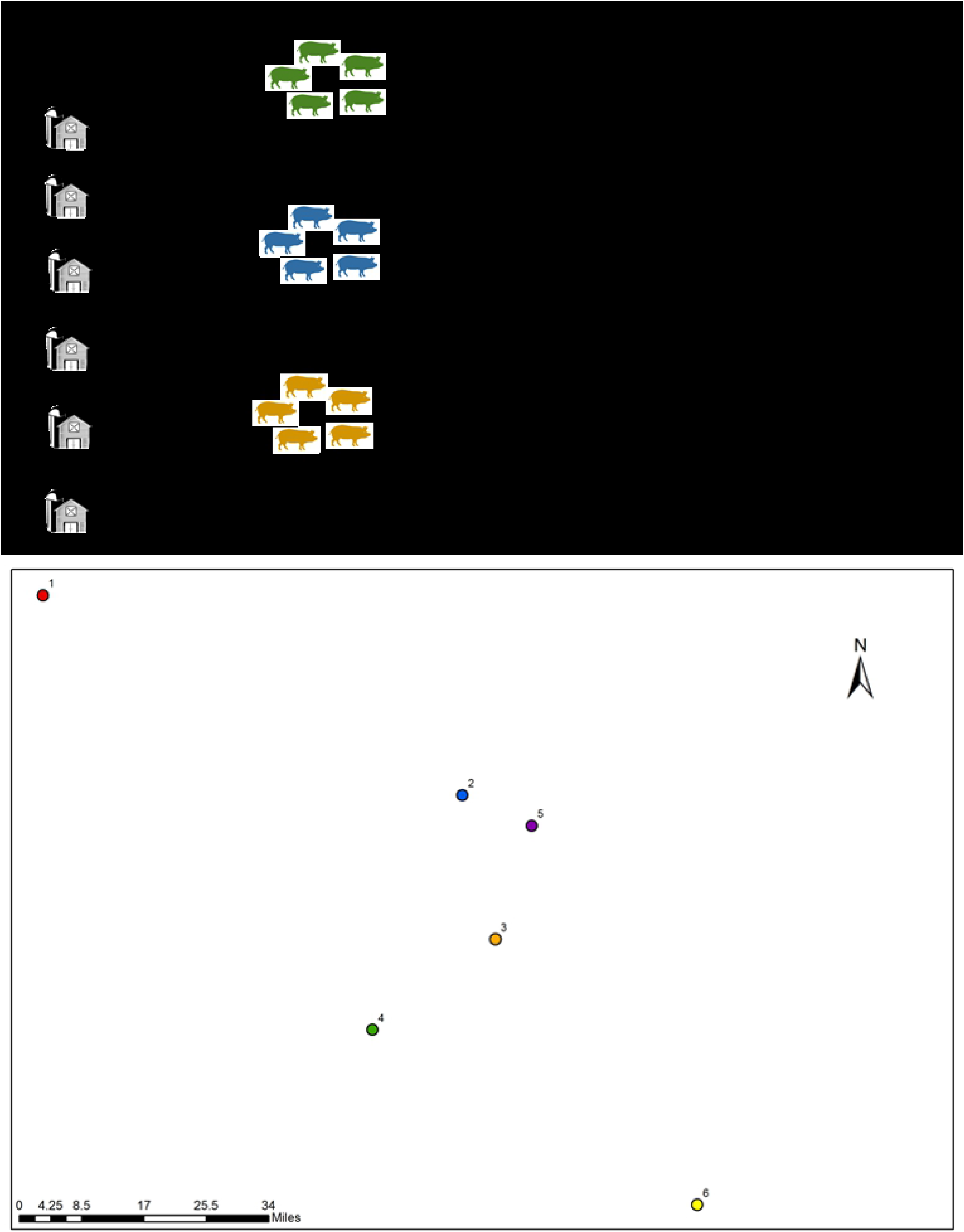
Sampling protocol showing study groups of interest and summarized sample processing steps; and relative locations of the six participating farms; with numbers representing farms 1-6. Actual map is not shown for confidentiality reasons.

### Sample pooling, genomic DNA extraction and sequencing

Fecal and nasal swab samples were transported to the laboratory located at the Food Animal Health Research Program (The Ohio State University) in Wooster, Ohio, in a sealed cooler with dry ice within 24 hours of collection. Two milliliters of phosphate-buffered saline (PBS) were added to each nasal swab dry tube and the samples were centrifuged at 13,000 g for 15 min. One milliliter of the supernatant was pelleted and resuspended into 250 μl of the original supernatant and the rest of the product was stored at −80°C for virome studies (unpublished data). Fecal and nasal swab samples were pooled across animals by farm and health status according to Fig. 1 (0.5 ± 0.015 g of feces per sample; 45 μl of nasal swab suspension per sample) in pools of five for a total of 72 pooled samples (36 fecal and 36 nasal swab pooled samples). Genomic DNA extraction of the pooled fecal (0.2 ± 0.015 g) or nasal (225 μl) swab samples was conducted using the PureLink Microbiome DNA Purification Kit (Life Technologies, Invitrogen Corp.) followed by RNAse treatment (10 units/hr), as previously described [12]. Next generation sequencing library preparations and Illumina MiSeq sequencing were conducted at the Bioscience Division, Los Alamos National Laboratory, NM. Briefly, DNA samples were quantified using a Qubit 2.0 Fluorometer (Invitrogen, Carlsbad, CA) and DNA quality was checked on a 0.6% agarose gel using E-gel electrophoresis system (ThermoFischer Scientific, Grand Island, NY). A total of 50 ng DNA was used to generate amplicons that cover V4-V5 hypervariable regions of bacteria and Archaea16S rRNA using F341-806R pair of primer. The first round of PCR amplified the 73-V4 region using KAPA HIFI HotStart Ready Mix (Kapa Biosystems, Wilmington, MA) with following PCR conditions; 95C for 3 minutes, 20 cycles of 95C for 30 seconds, 55C for 30 seconds and 72C for 30 seconds and an extension of 72C for 5 minutes. The second round of PCR added the Illumina specific sequencing adapter sequences and unique indexes using the Nextera XT Index Kit v2 (Illumina, San Diego, CA) and KAPA HIFI HotStart Ready Mix (Kapa Biosystems, Wilmington, MA) with following PCR conditions; 95C for 3 minutes, 8 cycles of 95C for 30 seconds, 55C for 30 seconds and 72C for 30 seconds and an extension of 72C for 5 minutes. The amplicons were cleaned up using AMPure XP beads (Beckman Coulter, Indianapolis, IN). The concentration of the amplicons pool was obtained using the Qubit dsDNA HS Assay (ThermoFisher Scientific, Grand Island, NY). The average size of the library was determined by the Agilent High Sensitivity DNA Kit (Agilent, Santa Clara, CA). An accurate library quantification was determined using the Library Quantification Kit – Illumina/Universal Kit (KAPA Biosystems, Wilmington, MA). The amplicon pool was sequenced on the Illumina MiSeq generating paired end 300 bp reads. The amplicon pool was demultiplexed using Illumina’s bcl2fastq.

### Taxonomic microbiota analyses and statistical analyses

Quality control and processing of the raw reads was performed using EDGE bioinformatic platform version 1.5.1, as previously described [13]. Annotation of the trimmed merged reads to the Greengenes reference database (08-2013 release) was performed using QIIME (quantitative insights into microbial ecology) software package version 1.9.1 with a similarity threshold of 94%. Alpha and beta diversity analyses were conducted using the core_diveristy_analysis.py script in QIIME. Samples with fewer than one hundred reads were removed from analyses. Observed OTUs, PD Whole Tree, and Chao1 were calculated in QIIME and compared according to sow health status (healthy, cull, compromised cull), and farm of origin (1-6) using Kruskal-Wallis test with Dunn’s post-hoc tests [14] in STATA/ version 14.2, College Texas, TX. Beta diversity was assessed using weighted UniFrac within QIIME and tested using PERMANOVA. Lastly, relative abundance data were analyzed using g-tests and p-values were FDR-corrected for multiple hypothesis testing (group_significance.py QIIME script). Samples were considered statistically different when the accompanying FDR-corrected p-value was < 0.05; and a tendency was described as p-value > 0.05 and < 0.10.

## Results

We obtained a total of 5,855,302 reads, with 1,715,767,300.5 total bases, and a mean read length of 293.03 base pairs. For 36 pooled fecal samples, we obtained 9,251 OTUs, and total count of 1,034,925 reads. For 35 pooled nasal samples (one sample was lost), we obtained 805 OTUs, and a total count of 134,609 reads. The mean number of OTUs per nasal pooled sample was 3,845.97 (SD: 9,078.87), the median was 1,069.00, with a minimum of 2.00, and a maximum of 52,465.00. One hundred OTUs was used as a cut-off, therefore, nine samples (out of 35) were not included in further analyses; and the final number used in the description below for nasal swab samples was 26. The mean number of OTUs per fecal pooled sample was 28,747.92, and the median was 26,410.00 (SD: 14,090.50), with a minimum of 6,004.0, and a maximum of 75,134.0. As expected, the diversity and richness of the nasal microbiota was low compared to the gut microbiota.

### Nasal microbial diversity and composition

There was no difference in alpha diversity indices between nasal pooled samples from compromised, cull, and healthy sows using either Chao1 (χ2(2) = 1.997, p = 0.361), observed OTUs (χ2(2) = 0.667, p = 0.716), or PD Whole Tree (χ2(2) = 1.985, p = 0.371; Table 2). Likewise, there was no difference in diversity indices between pooled samples from farms 1-6 using observed OTUs (χ2(5) = 8.039, p = 0.154), or PD Whole Tree (χ2(5) = 8.054, p = 0.1533); but there was a statistical difference when using Chao1 (χ2(5) = 22.311, p = 0.0005; Table 2; Fig 2). Furthermore, PERMANOVA on the PCoA based on weighted beta diversity values did not indicate clustering by farm (pseudo-F = 1.22, p = 0.22; Fig 3), but indicated clustering by health status (pseudo-F = 2.27, p = 0.017; Fig 3). In this case, PC1 explained approximately 42.78% of the variation, with samples obtained from healthy sows differentiating from the other two health status groups along this axis; PC2 accounted for approximately 24.63% of the variation with samples obtained from cull sows separating relatively clearly along this axis.

**Table 2.** Alpha diversity indices for nasal and fecal samples collected from sows in six farms and from different health status. Measures presented in means (SD), with letters representing statistical significant using Kruskal-Wallis test followed by the Dunn’s post-hoc test.

| Group | Nasal samples |  |  | Fecal samples |  |  |
| --- | --- | --- | --- | --- | --- | --- |
|  | PD Whole | Chao1 | Observed OTUs | PD Whole | Chao1 | Observed OTUs |
| All | 2.57<br>(0.76) | 41.31<br>(18.69) | 19.19<br>(7.58) | 36.08<br>(13.61) | 1037.63<br>(471.32) | 467.20<br>(225.25) |
| Farm 1 | 2.33<sup>a,b</sup><br>(0.59) | 32.56<br>(10.57) | 16.92<br>(5.72) | 35.06<br>(13.53) | 979.88<br>(460.09) | 453.98<br>(224.02) |
| Farm 2 | 2.23<sup>a,b</sup><br>(0.53) | 31.10<br>(10.25) | 15.58<br>(5.29) | 33.40<br>(12.87) | 962.70<br>(446.23) | 432.75<br>(214.49) |
| Farm 3 | 2.35<sup>a</sup><br>(0.59) | 29.53<br>(10.85) | 17.15<br>(5.52) | 33.50<br>(13.19) | 1029.64<br>(432.07) | 432.07<br>(220.05) |
| Farm 4 | 2.84<sup>b,c</sup><br>(0.86) | 41.89<br>(15.16) | 20.42<br>(7.77) | 37.84<br>(14.71) | 1083.49<br>(509.62) | 486.41<br>(242.39) |
| Farm 5 | 2.86<sup>c</sup><br>(0.99) | 56.80<br>(22.57) | 21.85<br>(9.26) | 38.00<br>(14.69) | 1092.38<br>(515.72) | 480.64<br>(242.30) |
| Farm 6 | 2.78<sup>c</sup><br>(0.79) | 55.98<br>(18.53) | 23.11<br>(9.19) | 38.69<br>(14.78) | 1077.69<br>(498.91) | 507.38<br>(249.15) |
| All | 2.61<br>(0.74) | 43.01<br>(14.99) | 19.73<br>(7.25) | 36.08<br>(13.55) | 1037.63<br>(472.56) | 467.20<br>(225.20) |
| Cull | 2.68<br>(0.81) | 47.14<br>(17.51) | 20.46<br>(7.87) | 34.82<br>(13.54) | 994.60<br>(469.87) | 448.35<br>(222.92) |
| Compromised | 2.75<br>(0.80) | 43.17<br>(14.10) | 20.15<br>(7.52) | 36.83<br>(14.19) | 1059.56<br>(493.18) | 480.48<br>(237.58) |
| Healthy | 2.37<br>(0.63) | 38.73<br>(13.19) | 18.57<br>(6.87) | 36.58<br>(14.15) | 1058.73<br>(497.54) | 472.78<br>(235.65) |

<sup>a,b,c</sup>Different letters represent statistically significant differences at $P < 0.05$

**Fig 2.**
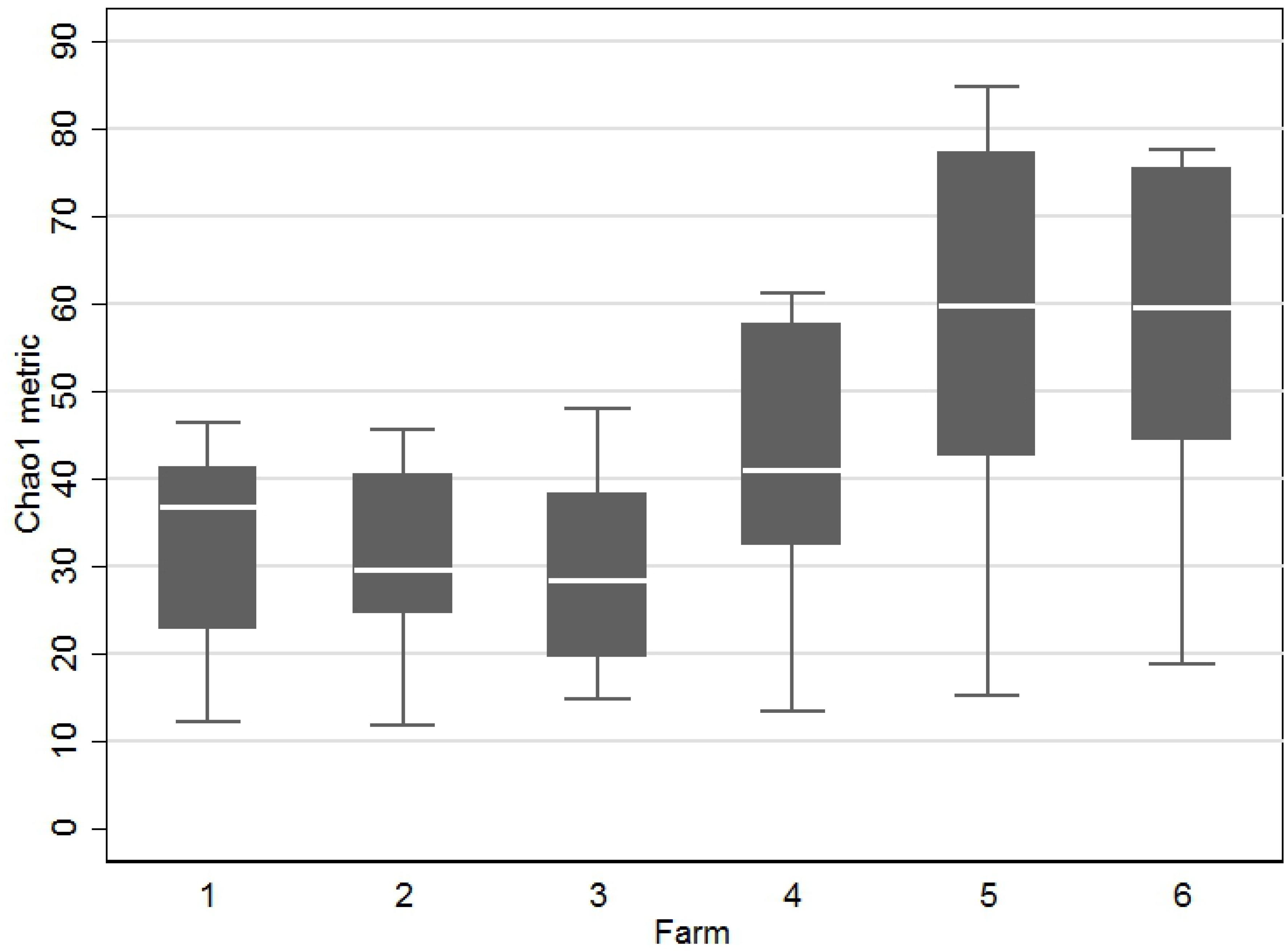
Boxplot showing alpha diversity represented by the Chao1 metric showing the difference in indices between nasal swab samples from animals from farms 1-6.

In terms of relative abundance, *Proteobacteria* (66.02 ± 12.86 [mean % ± standard deviation %]) was the largest represented phyla from the pooled nasal swab samples, followed by *Firmicutes* (16.97 ± 8.41), *Bacteroidetes* (7.22 ± 6.20) and *Actinobacteria* (8.52 ± 6.65; Fig 4). Within *Proteobacteria,* the most represented family was *Moraxellaceae* (61.28 ± 13.78). Globally, the second most abundant family found was *Weeksellaceae* (6.31 ± 3.20) from the *Bacteroidetes* phylum.

Furthermore, nasal microbiota analysis revealed that the pools analyzed shared many of the same nasal bacterial taxa. All analyzed samples (100%) shared four OTUs: phylum *Actinobacteria,* order *Actinomicetales*; phylum *Firmicutes,* orders *Lactobacillales* and *Clostridiales;* and plylum *Proteobacteria*, order *Pseudomonadales*. Common families of the shared *Actinobacter*ia OTUs included *Corynebacteriaceae*, and *Microccaceae*; common families of the shared *Firmicutes* OTUs included *Staphylococcaceae*, *Enterococcaceae*, and *Streptococcaceae*, and *Ruminococcaceae;* and common families of the shared *Proteobacteria* OTUs included *Neisseriaceae* and *Moracellaceae*.

Finally, in order to unravel the differences in nasal microbiota in sows from different health status and sows within different farms, we analyzed the OTU abundance at the genus level by grouping samples according to sow health status, and farm of origin. We found no statistically significant differences between the different groups of animals in regards to the nasal microbiota using the FDR-corrected p-values.

### Fecal microbial diversity and composition

There was no difference in diversity indices between fecal pooled samples from compromised, cull, and healthy sows using either Chao1 (χ2(2) = 0.286, p = 0.867), observed OTUs (χ2(2) = 0.329, p = 0.848), or PD Whole Tree (χ2(2) = 0.504, p = 0.777; Table 2). There was likewise no difference in diversity indices between fecal pooled samples from farms 1-6 using Chao1 (χ2(5) = 1.38, p = 0.927), observed OTUs (χ2(5) = 1.812, p = 0.874), or PD Whole Tree (χ2(5) = 3.874, p = 0.568; Table 2). PERMANOVA conducted on weighted PCoA showed an influence of farm of origin on the pooled fecal samples (pseudo-F = 4.66, p = 0.001; Fig 3). For all PCoA, PC1 explained approximately 9.45% of the variation, with samples along this axis separated roughly by farm of origin; with samples from farms 1, 2, and 3, separating from samples from farms 4, 5 and 6. PC2 accounted for approximately 5.04% of the variation. However, gut microbial communities did not appear to be clustered by health status (pseudo-F = 0.93, p = 0.51).

**Fig 3.**
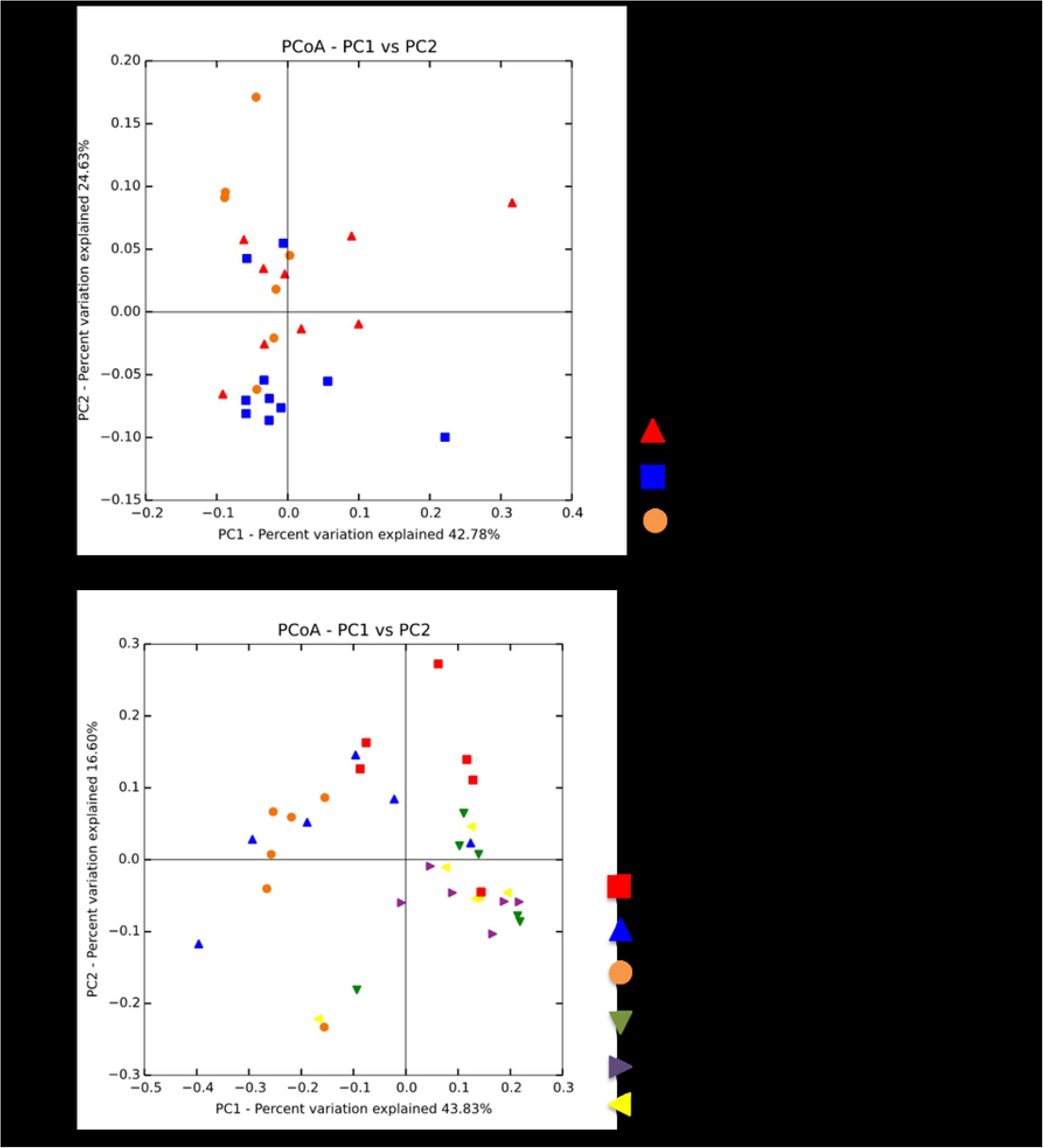
Principal component analysis (PCoA) plots generated using the weighted uniFrac from a) health status based on nasal swab samples, and b) farm of origin in fecal samples (different colors represent farms 1-6)

In terms of relative abundance, *Firmicutes* (61.93 ± 9.10) was the most represented phyla from the pooled fecal samples, followed by *Bacteroidetes* (15.52 ± 9.44), *Euryarchaeota* (11.20 ± 9.61), *Spirochaetes* (4.59 ± 3.42) and *Proteobacteria (2.18 ± 3.25)/Actinobacteria* (2.17 ± 0.83; Fig 4). Within *Firmicutes,* the most represented family was *Ruminococcaceae* (12.39 ± 4.13). Globally, the second most abundant family found was *Methanobacteriaceae* (11.17 ± 9.61) from the *Methanobacteria* phylum.

**Fig 4.**
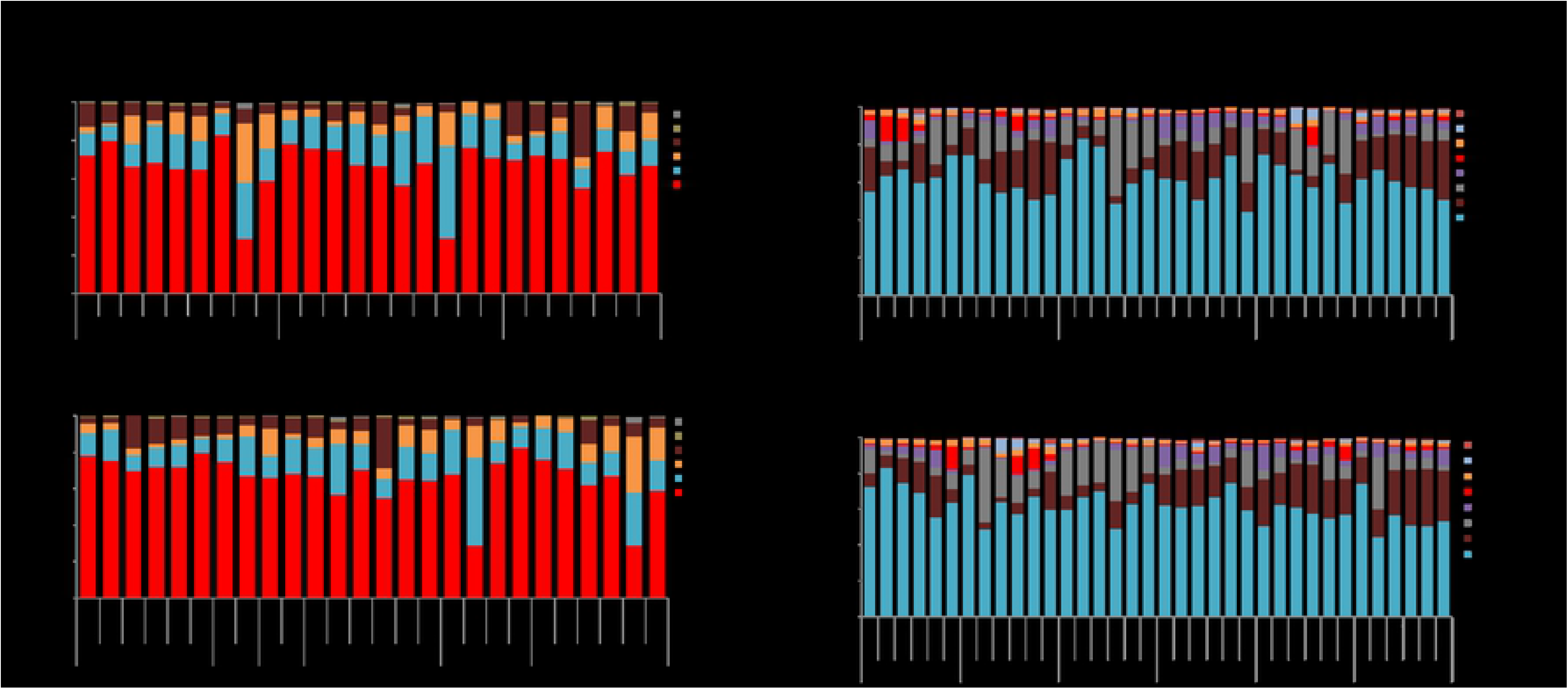
Distribution of relative abundance at phyla level for nasal swab (a) and fecal (b) samples; by health status and farm of origin. Phyla with relative abundance of less than 1% are not shown.

All analyzed samples shared various OTUs: phylum *Methanobacteria,* order *Methanobacteriales*; phylum *Actinobacteria*, orders *Bifidobacteriales, Coriobacteriales and Bacteroidales;* phylum *Firmicutes*, orders *Lactobacillales, Turicibacterales, Clostridiales,* and *Erysipelotrichales;* phylum *Proteobacteria,* order *Enterobacteriales;* and phylum *Spirochaetes*, order *Spirochaetales.* A common family of the shared *Methanobacteria* OTUs included *Methanobacteriaceae*; common families of the shared *Actinobacteria* OTUs included *Bifidobacteriaceae* and *Coriobacteriaceae;* common families of the shared *Bacteroidetes* OTUs included *Prevotellaceae, Paraprevotellaceae,* and *p-2534-18B5;* common families of the shared *Firmicutes* OTUs included *Lactobacillaceae, Streptococcaceae, Turicibacterraceae, Christensenellaceae, Clostridiaceae, Lachnospiraceae, Ruminococcaceae, Veillonellaceae, Mogibacteriaceae,* and *Erypelotrichaceae;* and common families from the OTUs *Proteobacteria* and *Spirochaetes* were *Enterobacteriaceae* and *Spirochaetaceae*, respectively.

Finally, significant differences at the genus level were found in the fecal microbiota composition of sows of different health status and farm of origin (Table 3). For health status, only one FDR-corrected value showed a trend towards significance (P = 0.0625), with compromised cull sows presenting a greater abundance (mean reads 943.82) of *Enterobacteriacea (Proteobacteria)* compared to cull, that had the lowest (mean reads 78.73) and healthy (mean reads: 358.64). In regards to differences among farms, nine genera/ families were identified as significantly different from at least one other group (Table 3). These were *Turicibacter, Bacteroidales, Clostridiaceae (SMB53 and Christensenellaceae), Bacteroidales (S24-7), Lachnospiraceae, Methanobrevibacter, Lactobacillus*, and *Treponema*.

**Table 3.** Results from analysis of relative abundance data; relative abundance was compared between farms and between health status for pooled sow fecal samples using g-tests and FDR-corrected p-value for multiple hypothesis testing. Mean values are shown; p-values < 0.05 were considered statistically significant and p-values between 0.05 and 0.10 were considered a tendency for significance.

| Taxonomy | Farm 1 | Farm 2 | Farm 3 | Farm 4 | Farm 5 | Farm 6 | FDR-corrected p-value |
| --- | --- | --- | --- | --- | --- | --- | --- |
| *Turicibacter* | 759.17 | 1773.40 | 1138.80 | 195.00 | 98.20 | 243.83 | 0.027 |
| *Bacterioidales* | 436.67 | 116.00 | 327.80 | 505.33 | 663.20 | 1640.83 | 0.027 |
| *SMB53 (Clostridiaceae)* | 356.17 | 929.20 | 800.20 | 144.67 | 47.80 | 200.00 | 0.031 |
| *Lachnospiraceae* | 17.67 | 184.20 | 28.80 | 1074.50 | 153.60 | 251.17 | 0.031 |
| *Methanobrevibacter* | 1694.83 | 679.60 | 636.60 | 1266.50 | 1190.60 | 2165.33 | 0.031 |
| *Lactobacillus* | 544.67 | 1605.60 | 2156.20 | 73.50 | 288.00 | 616.83 | 0.031 |
| *Treponema* | 5120.33 | 1326.40 | 2160.60 | 1435.83 | 1331.40 | 2799.50 | 0.033 |
| *Christensenellaceae* | 803.17 | 106.00 | 224.20 | 986.33 | 1245.60 | 1152.50 | 0.041 |
|  | Cull | Compromised | Healthy |  |  |  | FDR-corrected p-value |
| *Enterobacteriaceae* | 78.72 | 943.81 | 358.63 |  |  |  | 0.062 |

## Discussion

Cull sows are important from both a disease transmission and welfare perspective, but are an understudied subpopulation of animals within commercial swine farms. Studies looking at both gut and nasal microbiota of adult female pigs are lacking in the literature. To the knowledge of the authors, this study was the first to characterize gut and nasal microbiota for sows of different health status housed in six different commercial farms under field conditions. The main strength of our study is that we sampled a considerable number of animals from six different farms under commercial, “real-life” conditions and that were under management of the same swine production company, which minimized the effect of factors that are known to influence the microbiome such as host genetics, diet, biosecurity, vaccination and treatment protocols, besides others when comparing groups of interest.

Overall, the diversity of the nasal microbiota was low compared to the gut microbiota (Table 2), as have been previously reported in swine microbiota studies [15]. The mean and/ or median number of OTUs per nasal swab sample reported in our study is within the range previously reported in other swine studies (median of 6,257 OTUs per sample reported by Slifierz et al. [16]; and mean of 1,749 reported by Weese et al. [17]); and the median OTUs per fecal sample reported in our study is higher than previously reported (median of 1,976 OTUs per sample reported by Slifierz et al. [15]). We hypothesize that factors such as animal age, management, and laboratory methods (pooling versus individual samples, 16S hypervariable region, sequencing platform, etc.) may explain observed OTU difference in our study.

Our study showed differences in microbial diversity for nasal swab samples across animals from different farms; but no difference was found for nasal swab samples across animals from different health status; nor for any comparisons for fecal samples. Literature findings on swine nasal microbial diversity are few and conflicting; with significantly decreased diversity linked to clinical conditions such as Glasser’s disease [10]; but not to methicillin-resistance *Staphylococcus aureus* (MRSA) carriage [17]. Even though the farms enrolled in the current study were under the same management practice, had the same herd veterinarian, and overall health protocols, there are inherent differences regarding past disease challenges and herd productivity that were not captured in a reliable manner at this point. Future studies should focus on exploring the association between the nasal passage microbiota and detailed health and production parameters at the animal or herd levels.

Interestingly, PCoA diversity showed that health status had a stronger influence on the nasal microbial community composition than farm of origin. Findings differed when analyzing the fecal microbiota, where farm of origin was associated with distinct microbial community composition, with farms 1-3 and 4-6 clustered together. Interestingly, these clusters were not necessarily explained by geographical location, as seen on Fig 1. As previously discussed, even though in general farms from the same production system share many known factors, physical characteristics of the farm and other management-related variations (e.g. farm size, number of workers, environment, among others) may be implicated on the differences observed. In this case, smaller farm size and the fact that all animals were housed in individual crates were some differences observed when comparing farms 1-3 to 4-6 (Table 1). These may be proxy variables that were not investigated in this study but that could have an important impact on defining animal’s microbiota, such as number of workers and amount of within and outside-farm movements.

In terms of relative abundance in nasal swab samples, *Proteobacteria* was the largest represented phyla, followed by *Firmicutes, Bacteroidetes* and *Actinobacteria*. These results are similar to swine-related nasal microbiota studies conducted in Canada, the United Kingdom and Spain [10,16-17] which report *Proteobacteria* as the most abundant phyla, followed by a large abundance in *Bacteroidetes*, and *Firmicutes*. In contrast, the abundance of *Actinobacteria* in our study is higher than what was previously reported [10,16], and the abundance of *Tenericutes* (particularly *Mycoplasmacetae* family) was more prominent in the European study [10] compared to our current study. These differences could be related to geography, animal management and age (both mentioned studies were conducted with piglets 4-6 weeks of age or slaughter animals [approximately 6 months of age], which would be younger than our adult female study population), laboratory methods, besides other factors. In regards to beta diversity, Correa-Fiz et al. [10] reported significant nasal microbiota differences by health status; similarly to what we described in our current study. However, it is important to note that health status was determined at the farm level for that study, and therefore, these two were confounded and their separate effects could not be definitively characterized.

Most of the shared taxa for fecal samples in our study were in the *Firmicutes* and *Bacteroidetes* families. This agrees with a meta-analysis published in 2017, which reported that those two phyla accounted for nearly 85% of the 16S rRNA gene sequences among over 930 swine gastrointestinal samples [18]. This meta-analysis also points out that the main influencing factors of swine gut microbiota were study itself, GI location, and animal age; which makes comparisons of findings between studies a challenge. A study by Kim et al. [15] analyzed fecal samples for pigs of various ages (including sows) and reported that the most common microbial families from fecal samples from the five sampled sows in such study were *Prevotellaceae*, *Ruminococcaceae, Lachnospiraceae*, and *Streptococcaceae*, which were also present and commonly shared among pools in our study. Those authors also report a large proportion of “unclassified” species, which did not occur in our case.

Numerous genera/ families were over- or under-represented between samples coming from different farms; and the *Enterobacteriaceae* family was over-represented in compromised animals and under-represented in cull animals, compared to healthy animals (Table 3). Due to the lack of farm-level information on performance or health concerns, farm-level microbial comparisons as it relates to health or production cannot be made at this point. The *Enterobacteriaceae* family is known to be composed of important commensal and potentially pathogenic microbes [19]. Our findings did not allow for genera differentiation, but we hypothesize that cull animals are immunocompromised and therefore; would have a less diverse *Enterobacteriaceae* presence of commensal microbiota; on the other hand, compromised animals (which would be animals identified by the acute nature of their condition on farm) may have an increase of potentially pathogenic microbes, including those of the *Enterobacteriaceae* family. This study had limitations. Firstly, the samples analyzed here were pools of five pigs. Even though these animals were pooled by our main variable of interest; health status; this resulted in loss of potential for analysis of individual-level variables such as parity, age, and reproductive performance. Furthermore, this also likely had an impact in microbe abundance and richness. Another limitation is that, even though farms were under the same management system and protocols, data in regards to past farm health challenges and actual antibiotic usage for the sampled animals is unknown. During informal discussions with farm managers in relation to this cross-sectional study, information provided was that none of the herds had gone through health challenges of note (e.g., considerable disease outbreaks or increase in antimicrobial use for specific prevention or treatment reasons); but this was not systematically captured retrospectively by the investigators. We further acknowledge the potential for misclassification bias when selecting study subjects particularly for the cull and healthy sows; since investigators relied on sow cards for identification of culling animals; but confirmation with the farm manager was attempted whenever possible to minimize this potential source of bias.

Future studies should focus on expanding sample size, considering individual samples, and capturing information such as animal age, antimicrobial treatment, and production records so that these important variables could be taken into consideration during the analytical phases.

## Conclusions

In conclusion, this study provided baseline information for nasal and fecal microbiota of sows under field conditions. Our results suggest that farms within the same production system and different health status can affect microbial diversity and composition for such examined body regions.

## Acknowledgements

The authors would like to acknowledge farm owners, managers and personnel, and undergraduate students for help with sampling collection.

## References

1. D’Allaire S, Stein TE, Leman AD. Culling patterns in selected Minnesota swine breeding herds. 1987. Can. J. Vet. Res. 1987; 51:506–512.

2. Stalder KJ, Lacy RC, Cross TL, Conatser GE. Financial impact of average parity of culled females in a breed-to-wean swine JSHAP. 2003; 11: 69–74.

3. Stein TE, Dijkhuizen A, D’Allaire S, Morris RS. Sow culling and mortality in commercial swine breeding herds. Prev. Vet. Med. 1990; 9: 85–94. https://doi.org/10.1016/0167-5877(90)90027-F

4. Sutherland, D. The marketing journey of cull sows and secondary market pigs. Swine Health Information Center. 2017. https://www.swinehealth.org/wp-content/uploads/2018/07/The-Marketing-Journey-of-Cull-Sows-and-Secondary-Market-Pigs.pdf. Last accessed March 14th, 2019.

5. Round JL, Mazmanian SK. The gut microbiota shapes intestinal immune responses during health and disease. Nat. Rev. Immunol. 2009; 9:313–323. https://doi.org/10.1038/nri2515

6. Young VB. The role of the microbiome in human health and disease: an introduction for clinicians. BMJ 2017; 356, j831. https://doi.org/10.1136/bmj.j831

7. Nowland TL, Plush KJ, Barton M, Kirkwood RN. Development and Function of the Intestinal Microbiome and Potential Implications for Pig Production. Animals. 2019. https://doi.org/10.3390/ani9030076

8. Jami E, White BA, Mizrahi I. Potential Role of the Bovine Rumen Microbiome in Modulating Milk Composition and Feed Efficiency. PLOS ONE 2014; 9:e85423. https://doi.org/10.1371/journal.pone.0085423

9. Niederwerder MC. Role of the microbiome in swine respiratory disease. Alternative strategies for the control of porcine reproductive and respiratory syndrome. Vet. Microbiol. 2017; 209:97–106. https://doi.org/10.1016/j.vetmic.2017.02.017

10. Correa-Fiz F, Fraile L, Aragon V. Piglet nasal microbiota at weaning may influence the development of Glässer’s disease during the rearing period. BMC Genom. 2016; 17. https://doi.org/10.1186/s12864-016-2700-8

11. Niederwerder MC, Jaing CJ, Thissen JB, Cino-Ozuna AG, McLoughlin KS, Rowland RRR. Microbiome associations in pigs with the best and worst clinical outcomes following co-infection with porcine reproductive and respiratory syndrome virus (PRRSV) and porcine circovirus type 2 (PCV2). Vet. Microbiol. 2016; 188:1–11. https://doi.org/10.1016/j.vetmic.2016.03.008

12. Deblais L, Helmy YA, Kathayat D, Huang H-C, Miller SA, Rajashekara G. Novel Imidazole and Methoxybenzylamine Growth Inhibitors Affecting Salmonella Cell Envelope Integrity and its Persistence in Chickens. Sci. Rep. 2018; 8:13381. https://www.nature.com/articles/s41598-018-31249-0. Last accessed 01/22/2019.

13. Li PE, Lo CC, Anderson JJ, Davenport KW, Bishop-Lilly KA, Xu Y, Ahmed S, Feng S, Mokashi VP, Chain PSG. Enabling the democratization of the genomics revolution with a fully integrated web-based bioinformatics platform. Nucleic Acids Res. 2017; 45: 67–80. https://doi.org/10.1093/nar/gkw1027

14. Dinno A. Dunntest: a Stata package to perform Dunn’s pairwise multiple tests. 2014. https://alexisdinno.com/stata/dunntest.html.

15. Kim HB, Isaacson R. The pig gut microbial diversity: understanding the pig gut microbial ecology through the next generation high throughput sequencing. Vet. Microbiol. 2015; 177: 242–251.

16. Slifierz MJ, Friendship RM, Weese JS. Longitudinal study of the early-life fecal and nasal microbiotas of the domestic pigs. BMC Microbiol. 2015; 15:184.

17. Weese JS, Slifierz M, Jalali M, Friendship R. Evaluation of the nasal microbiota in slaughter-age pigs and the impact on nasal methicillin-resistant Staphylococcus aureus (MRSA) carriage. BMC Vet. Res. 2014; 10:69.

18. Holman DB, Brunelle BW, Trachsel J, Allen HK. Meta-analysis to define a core microbiota in the swine gut. mSystems 2017; 2:e00004–17. http://doi.org/10.1128/mSystems.00004-17.

19. Schierack P, Walk N, Reiter K, Weyrauch KD, Wieler LH. Composition of *Enterobacteriaceae* populations of healthy domestic pigs. Microbiol. 2007; 153:3830–3837. doi: 10.1099/mic.0.2007/010173-0.

